# Sources of genomic diversity in the self-fertile plant pathogen, *Sclerotinia sclerotiorum*, and consequences for resistance breeding

**DOI:** 10.1101/2020.12.03.409698

**Authors:** Lone Buchwaldt, Harsh Garg, Krishna D. Puri, Jonathan Durkin, Jennifer Adam, Myrtle Harrington, Debora Liabeuf, Alan Davies, Dwayne D. Hegedus, Andrew G. Sharpe, Krishna Kishore Gali

**Affiliations:** Agriculture and Agri-Food Canada, Saskatoon Research and Development Centre, 107 Science Place, Saskatoon, Canada, S7N 0X2. School of Life and Environmental Sciences, the University of Sydney, NSW, Australia.; Department of Plant Pathology, University of California, Davis, CA, USA.; Driscoll’s, Watsonville, California, USA.; Global Institute for Food Security, University of Saskatchewan, 110 Gymnasium Place, Saskatoon, SK, Canada, S7N 0W9.; Crop Development Centre, Department of Plant Sciences, University of Saskatchewan, SK, Canada, S7N 5A8.

**Author notes:** These authors contributed equally to the manuscript (Harsh Garg, Krishna D. Puri).

**Keywords:** *Sclerotinia sclerotiorum*, *Brassica napus*, canola, mycelium compatibility, clones, SSR markers, population structure, outcrossing, aggressiveness, quantitative resistance

## Abstract

The ascomycete, *Sclerotinia sclerotiorum*, has a broad host range and causes yield loss in dicotyledonous crops world wide. Genomic diversity and aggressiveness were determined in a population of 127 isolates from individual canola (*Brassica napus*) fields in western Canada. Genotyping with 39 simple sequence repeat (SSR) markers revealed each isolate was an unique haplotype. Analysis of molecular variation showed 97% was due to isolate and 3% to geographical location. Testing of mycelium compatibility identified clones of mutually compatible isolates, and stings of pairwise compatible isolates not seen before. Importantly, mutually compatible isolates had similar SSR haplotype, in contrast to high diversity among incompatible isolates. Isolates from the Province of Manitoba had higher allelic richness and higher mycelium compatibility (61%) than Alberta (35%) and Saskatchewan (39%). All compatible Manitoba isolates were interconnected in clones and strings, which can be explained by wetter growing seasons and more susceptible crops species both favouring more mycelium interaction and life cycles. Analysis of linkage disequilibrium rejected random recombination, consistent with a self-fertile fungus and restricted outcrossing due to mycelium incompatibility, and only one meiosis per lifecycle. More probable sources of genomic diversity is slippage during DNA replication and point mutation affecting single nucleotides, not withstanding the high mutation rate of SSRs compared to genes. It seems accumulation of these polymorphisms lead to increasing mycelium incompatibility in a population over time. A phylogenetic tree grouped isolates into 17 sub-populations. Aggressiveness was tested by inoculating one isolate from each sub-population onto *B. napus* lines with quantitative resistance. Results were significant for isolate, line, and isolate by line interaction. These isolates represent the genomic and pathogenic diversity in western Canada, and are suitable for resistance screening in canola breeding programs. Since the *S. sclerotiorum* life cycle is universal, conclusions on sources of genomic diversity extrapolates to populations in other geographical areas and host crops.

**Author summary:** *Sclerotinia sclerotiorum* populations from various plant species and geographical areas have been studied extensively using mycelium compatibility tests and genotyping with a shared set of 6-13 SSR markers published in 2001. Most conclude the pathogen is clonally propagated with some degree of outcrossing. In the present study, a population of *S. sclerotiorum* isolates from 1.5 million km^2^ area in western Canada were tested for mycelium compatibility, and genotyped with 9 published and 30 newly developed SSR markers targeting all chromosomes in the dikaryot genome (8+8). A new way of visualizing mycelium compatibility results revealed clones of mutual compatible isolates, as well as long and short strings of pairwise compatible isolates. Importantly, clonal isolates had similar SSR haplotype, while incompatible isolates were highly dissimilar; a relationship difficult to discern previously. Analysis of population structure found a lack of linkage disequilibrium ruling out random recombination. Outcrossing, a result of alignment of non-sister chromosomes during meiosis, is unlikely in *S. sclerotiorum*, since mycelium incompatibility prevents karyogamy, and compatibility only occur between isolates with similar genomic composition. Instead, genomic diversity comprise transfer of nuclei through hyphal anastomosis, allelic modifications during cell division and point mutation. Genomic polymorphisms accumulate over time likely result in gradual divergence of individuals, which seems to resemble the ‘ring-species’ concept. We are currently studying whether nuclei in microconidia might also contribute to diversity. A phylogenetic analysis grouped isolates into 17 sub-populations. One isolate from each sub-population showed different level of aggressiveness when inoculated onto *B. napus* lines previously determined to have quantitative resistance to a single isolate. Seed of these lines and *S. sclerotiorum* isolates have been transferred to plant breeders, and can be requested from the corresponding author for breeding purposes. Quantitative resistance is likely to hold up over time, since the rate of genomic change is relatively slow in *S. sclerotiorum*.

## Introduction

The ascomycete plant pathogen, *Sclerotinia sclerotiorum*, survives in the soil for several years as sclerotia (resting bodies) consisting of condensed hyphae surrounded by a melanised rind. In the spring, sclerotia in the top soil layer germinated with apothecia containing ascospores that are dispersed by wind to surrounding plants. Ascospores are unable to penetrate the plant epidermis directly. Instead, they germinate with hyphae that colonize dead organic material and form infection cushions [1]. These provide energy for production of virulence factors, which enable the pathogen to penetrated the plant epidermis. The fungal hyphae colonize plants without forming secondary spores, thus, the pathogen only has a single spore generation per cropping season.

*Sclerotinia sclerotiorum* is a dikaryot with two nuclei in cells of actively growing hyphal tips and in each ascospore resulting from one meiotic and one mitotic cell division. In addition, a single nucleus is found in microconidia, but their role in the lifecycle has not been determined. Hyphal tips from two different isolates can unite in anastomosis as first demonstrated by Kohn et al. [2]. Later, Ford et al. [3] observed transfer of nuclei between anastomosed hyphae using auxotrophic mutants to validate the formation of new dikaryons with restored nutrient requirements. Paring of two *S. sclerotiorum* isolates on nutrient agar can distinguish between mycelium compatible isolates that grow as one colony, and isolates that are incompatible growing as two colonies separated by a line of sparse mycelium. All isolates are self-compatible when paired with itself. Test of mycelium compatibility can be applied to an entire population using a isolate by isolate paring matrix. Populations studies comprising *S. sclerotiorum* isolates from various plant species and geographical areas have used a combination of mycelium compatibility test, genotyping with molecular markers, and tests for aggressiveness on selected host lines. A set of simple sequence repeat (SSR) markers was published in 2001 [4], and has been widely used to genotype *S. sclerotiorum* isolates. Different levels aggressiveness among isolates inoculated onto various host species have been demonstrated [5, 6, 7]. A summary of results from numerous studies were reviewed by Petrofeza et al. [8]. Most studies agree *S. sclerotiorum* is clonally propagated with some level of outcrossing. Whether there is a relationship between isolates belonging to the same mycelium compatible group and their SSR haplotype has been difficult to discern.

The pathogen has a wide host range among dicotyledonous plant species including canola (*Brassica napus*), bean (*Phaselous vulgaris*), soybean (*Glycine max*), lentil (*Lens culinaris*) and sunflower (*Helianthus annus*) [9]. Each year around eight million hectares are planted to canola in the Canadian provinces of Alberta (AB), Saskatchewan (SK) and Manitoba (MB). A high earning potential for canola seed has led to shortening of crop rotations, which in turn has resulted in higher disease pressure from *S. sclerotiorum* that causes stem rot also know as white mould. Infection of canola occur during flowering when the spores grow on fallen petals and pollen adhering to plant surfaces. The most severe yield loss results from colonisation of the main stem which restricts vascular transport of water and nutrients to the seed. Stem symptoms consists of long, pale lesions that initially have a dark margin between infected and healthy tissues. Later, the lesions expand leading to soft and collapsed stems.

Resistance to *S. sclerotiorum* in *B. napus* as well as other crop species is a quantitative trait that rely on several defense genes and pathways with cumulative effect [7, 10]. Previously, we identified *B. napus* lines with quantitative resistance after screening of more than 400 germplasm lines obtained from gene banks worldwide [11]. The phenotyping method involved attaching mycelium plugs of a pathogen isolate to the main stem of flowering plants, thereby resembling the natural infection process described above. Germplasm lines were ranked from resistant to susceptible based on lesion length and percent soft and collapsed stems. Heterogeneity in the disease phenotype was eliminated in a sub-set of lines by repeated cycles of inoculation, selection of resistant plants, selfing and re-testing, and resulted in high level of quantitative resistance in four lines, PAK54 and PAK93 originating from Pakistan, DC21 from South Korea and K22 from Japan. The lines were phenotyped for disease reaction using a single *S. sclerotiorum* isolate, 321, collected in 1992 from a canola field in Olds, Alberta [12].

The present research seeks to characterize the genomic and pathogenic diversity in a *S. sclerotiorum* population from commercial canola fields throughout a large production area of western Canada measuring approximately 1500 kilometers from East to West and 1000 kilometers from North to South. The results have direct application for development of resistant canola varieties, by providing selected isolates that represent the diversity needed for screening of breeding lines. Importantly, the study identifies mechanisms leading to this diversity, and assesses the rate of change in *S. sclerotiorum* populations applicable across geographical areas and host species.

## Results

### Disease incidence and isolate collection

A survey of 168 commercial canola fields in 2010 for the incidence of *S. sclerotiorum* showed the pathogen was present in 88% (Fig 1). In Manitoba, the pathogen was found in all fields with 13% of fields having between 6 and 30% disease incidence (Table 1). In Saskatchewan, 49% of fields had no or traces of the disease on side branches and pods, while only 4% of fields had incidences between 6 to 30. In Alberta, all fields had only up to 5% disease incidence. Where possible four isolations were made from 10 plants per field. The total collection consisted of 1392 individual *S. sclerotiorum* isolates kept as sclerotia in long-term storage under cold and dry conditions.

**Table 1.**
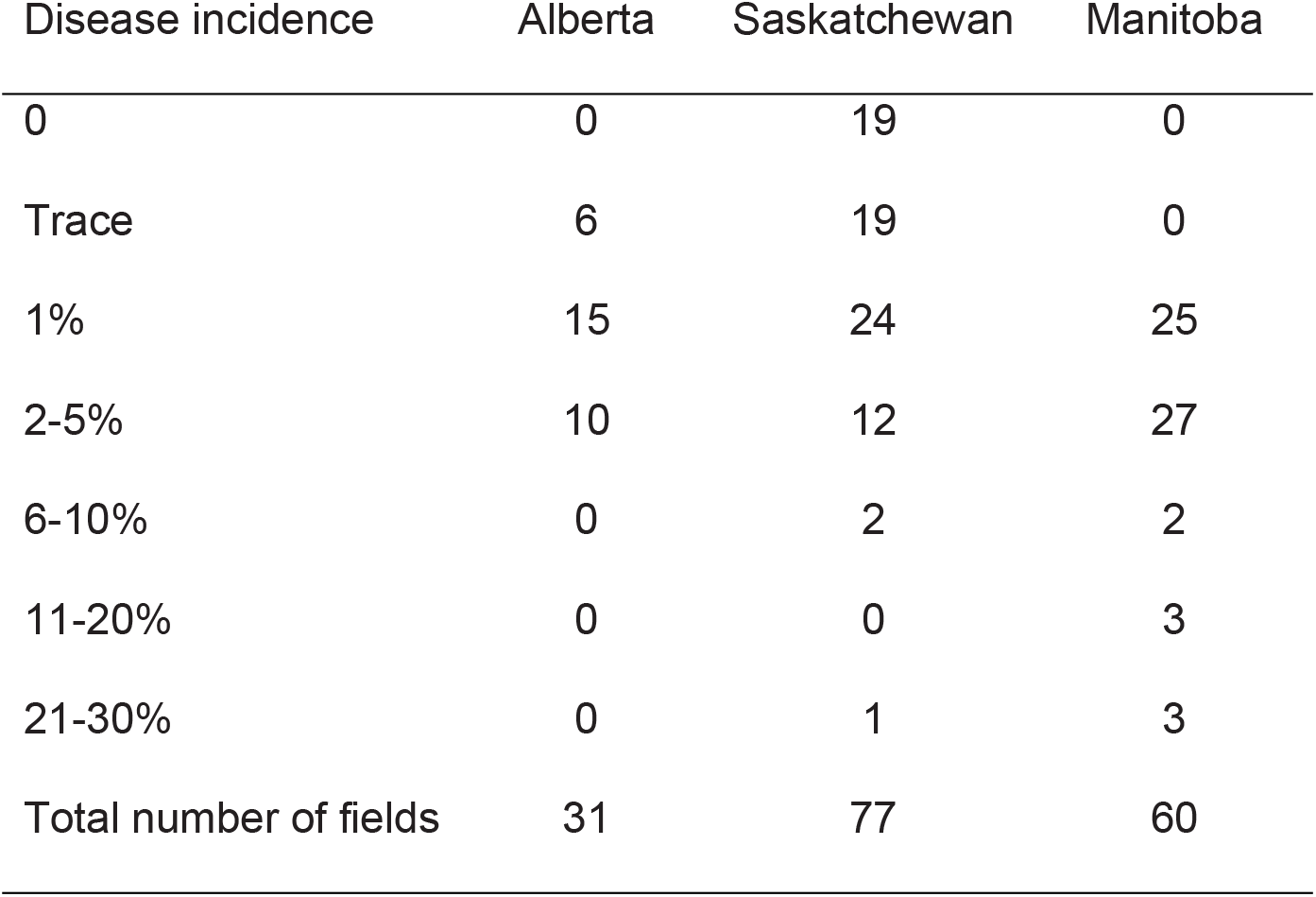
Incidence of *S. sclerotiorum* in commercial canola (*B. napus*) fields in three Canadian provinces.

| Disease incidence | Alberta | Saskatchewan | Manitoba |
| --- | --- | --- | --- |
| 0 | 0 | 19 | 0 |
| Trace | 6 | 19 | 0 |
| 1% | 15 | 24 | 25 |
| 2-5% | 10 | 12 | 27 |
| 6-10% | 0 | 2 | 2 |
| 11-20% | 0 | 0 | 3 |
| 21-30% | 0 | 1 | 3 |
| Total number of fields | 31 | 77 | 60 |

**Fig 1.**
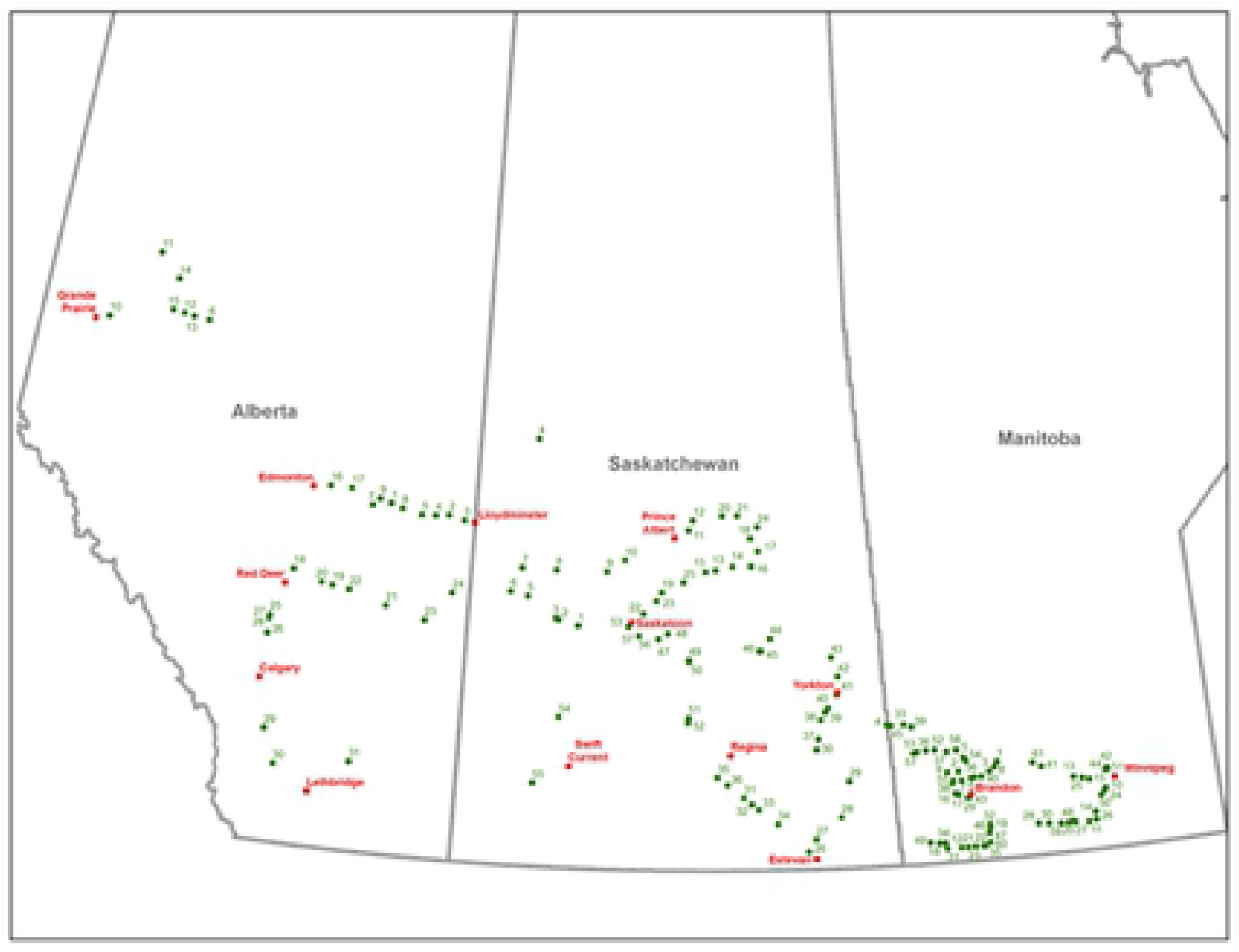
Geographical locations of commercial canola fields where *S. sclerotiorum* isolates were collected. A disease survey and isolate collection was carried out in 2010 across three Canadian Provinces. The sites closely resemble the distribution of canola producing areas (S6 Figure). A single isolate, 321, was collected in 1992.

### Mycelium compatibility relationships

A total of 133 *S. sclerotiorum* isolates from individual canola fields, as well as 36 isolates from a single heavily infected field in Saskatchewan, were tested for mycelium compatibility on PDA media in a replicated test for a total of 7734 isolate parings. Each isolate paired with itself in all cases by forming a single colony, thus confirming they were self-compatible. Isolates that were compatible with at least one other isolate within the same province comprised 61% in Manitoba, 39% in Alberta and 35% in Saskatchewan. For comparison, 41% were compatible within a single Saskatchewan field, while only 11% of isolates from different provinces were compatible (S1 Table).

Diagrams were created to visualize the relationship among compatible isolates in each isolate x isolate paring matrix by province, inter-province and in a single field as shown in Figs 2 and 3 (S1 Table). Circles specify groups of mutually compatible isolates belonging to the same clone, while arrows specify compatibility between pairs of isolates. Some isolates formed short and long strings where neighbouring isolates were compatible, but other isolates along the string were incompatible. Interestingly, all compatible isolates from Manitoba were closely related either as members of clones (A, B and C) or strings (Fig 2). Some of the longest strings consisted of 10 isolates, such as MB27-MB26-MB51-MB22-MB24-MB49-MB30-MB6-MB38-MB19. Compatible isolates in Saskatchewan were not as closely related forming only one clone and six pairs of compatible isolates, but no strings (Fig 3). It is striking that mycelium compatibility among isolates from 18 different canola fields in Saskatchewan closely resembled compatibility of 15 isolates from a single field in the same province, both having a single clone and five or six pairs of compatible isolates (Fig 3). Isolates in Alberta were not closely related, since neither clones nor strings were identified. When isolates from different provinces were paired, only the older isolate, 321, collected in 1992 from Alberta, was compatible with MB29 from Manitoba.

**Fig 2.**
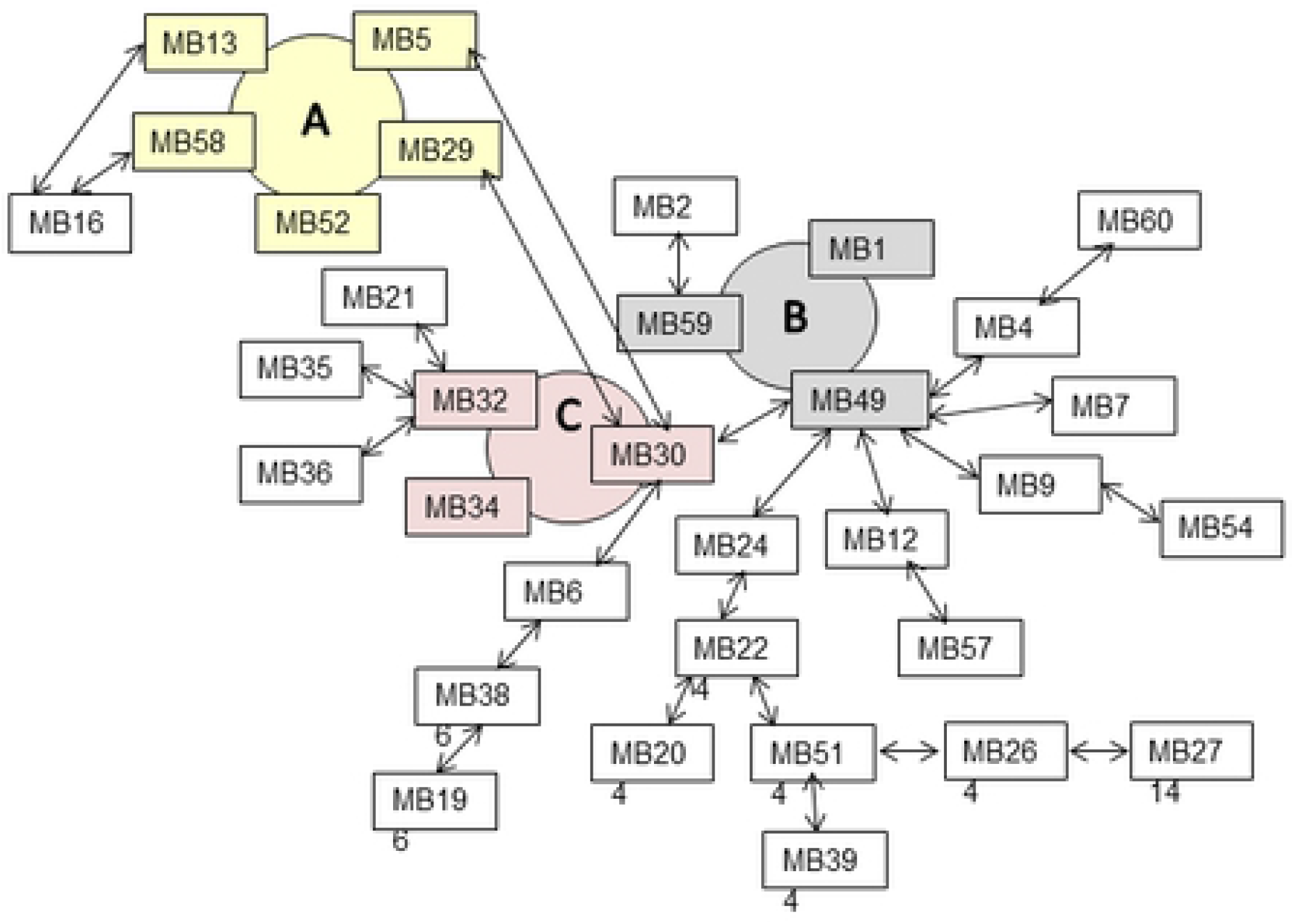
Mycelium compatibility among *S. sclerotiorum* isolates from canola fields in Manitoba. Coloured circles show mutually compatible isolates, and arrows show compatibility between two isolates. Table S1 contains all isolate by isolate scoring matrixes.

**Fig 3.**
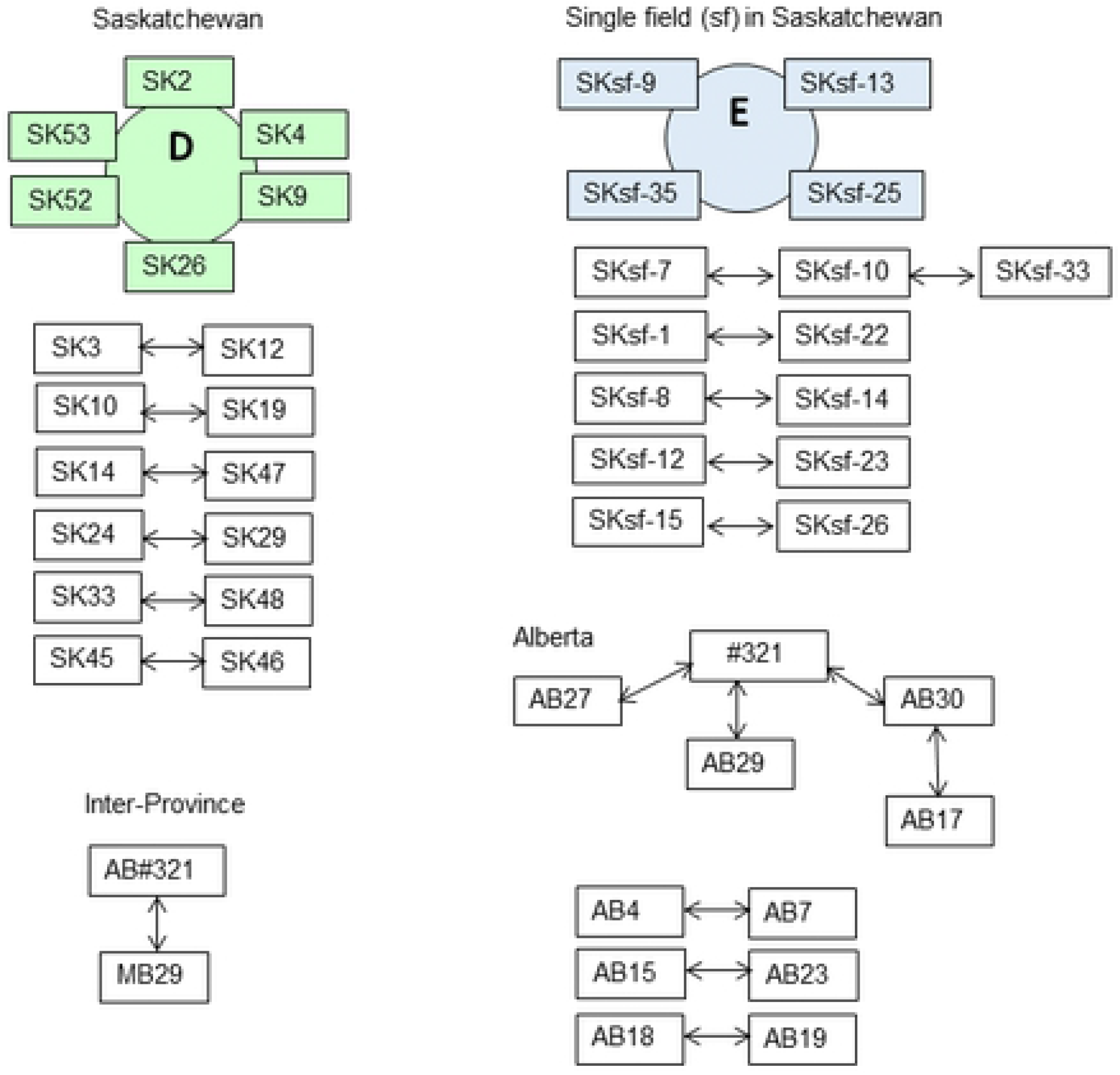
Mycelium compatibility among *S. sclerotiorum* isolates from canola fields in Saskatchewan, Alberta, a single field, and inter-Province. Coloured circles show clones of mutually compatible isolates, and arrows show compatibility between two isolates. Table S1 contains all isolate by isolate scoring matrixes.

### Simple sequence repeat polymorphisms

The sequenced *S. sclerotiorum* genome [13] was used to design 37 primer pairs for amplification of simple sequence repeats (SSR), so that 15 chromosomes and one contig were represented (labelled AAFC). Another 15 SSR were selected from the literature, (labelled ssr) [4]. A subset of 39 SSR (30 AAFC and 9 ssr) provided high quality amplification products, and were used to genotype 127 *S. sclerotiorum* isolates collected in individual fields (Table 2). The resulting scoring matrix consisted of 396 polymorphic alleles. Each SSR primer pair amplified between 2 and 35 alleles, of which 75% were shared by two or more isolates, while the remaining 25% were private alleles present in only one isolate. The values for polymorphic information content (PIC) and genomic diversity (*H*) for each SSR marker were highly correlated (r = 0.99) and ranged from 0.126 to 0.949 and 0.136 to 0.959, respectively (Table 2). The PIC value was above 0.5 for 81% (26) of the AAFC markers. The following markers on 12 chromosomes were particularly informative since they each amplified a highly polymorphic locus shared by many isolates: AAFC-2d on chromosome 1, AAFC-22c on chromosome 2, AAFC-3c on chromosome 4, AAFC-7b on chromosome 5, AAFC-9b on chromosome 6, AAFC-6f on chromosome 7, AAFC-11a on chromosome 9, AAFC-20b on chromosome 11, AAFC-33d on chromosome 12, AAFC-12a on chromosome 13, AAFC-25d on chromosome 14, AAFC-15e on chromosome 15, and AAFC-4d on contig R.

**Table 2.**
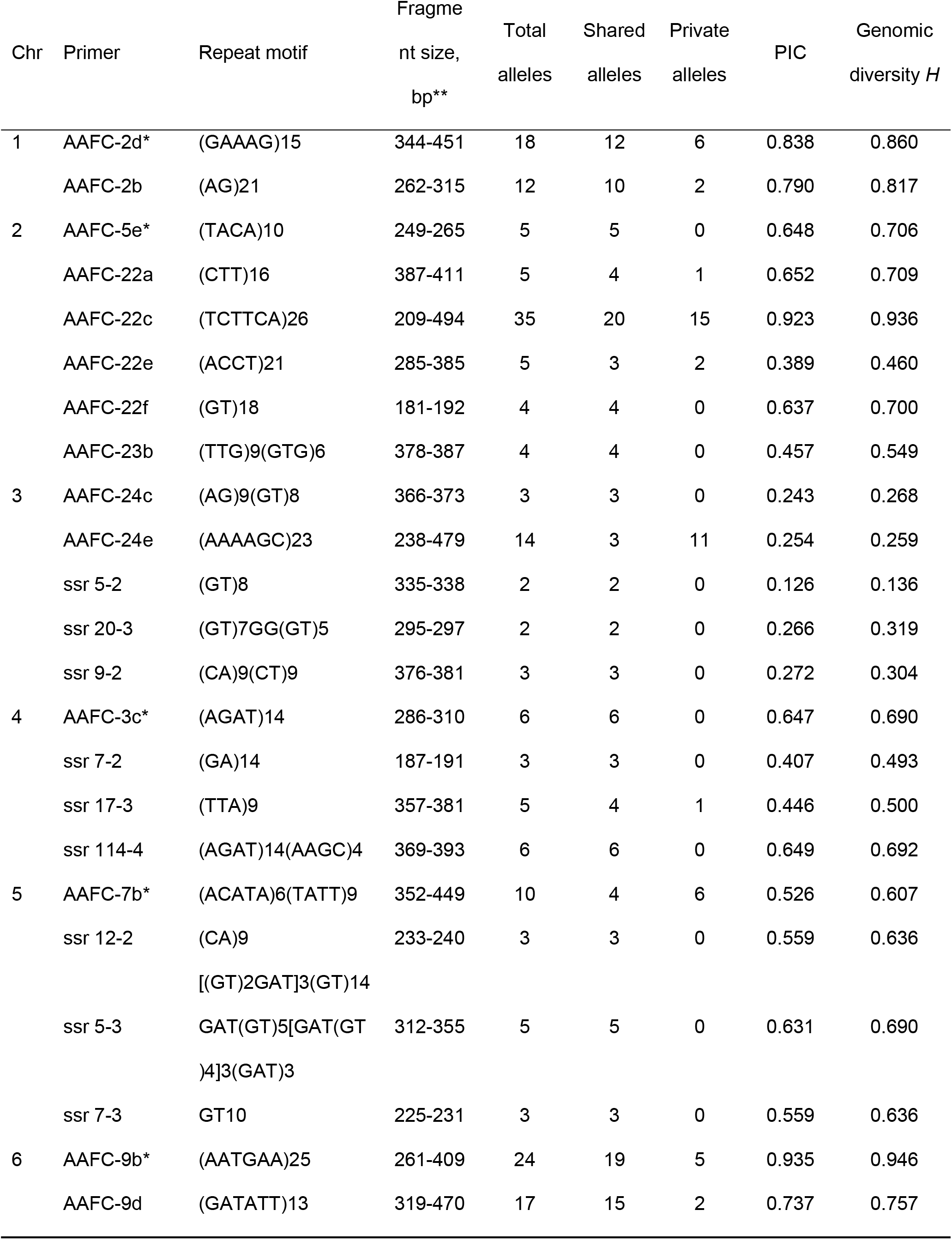
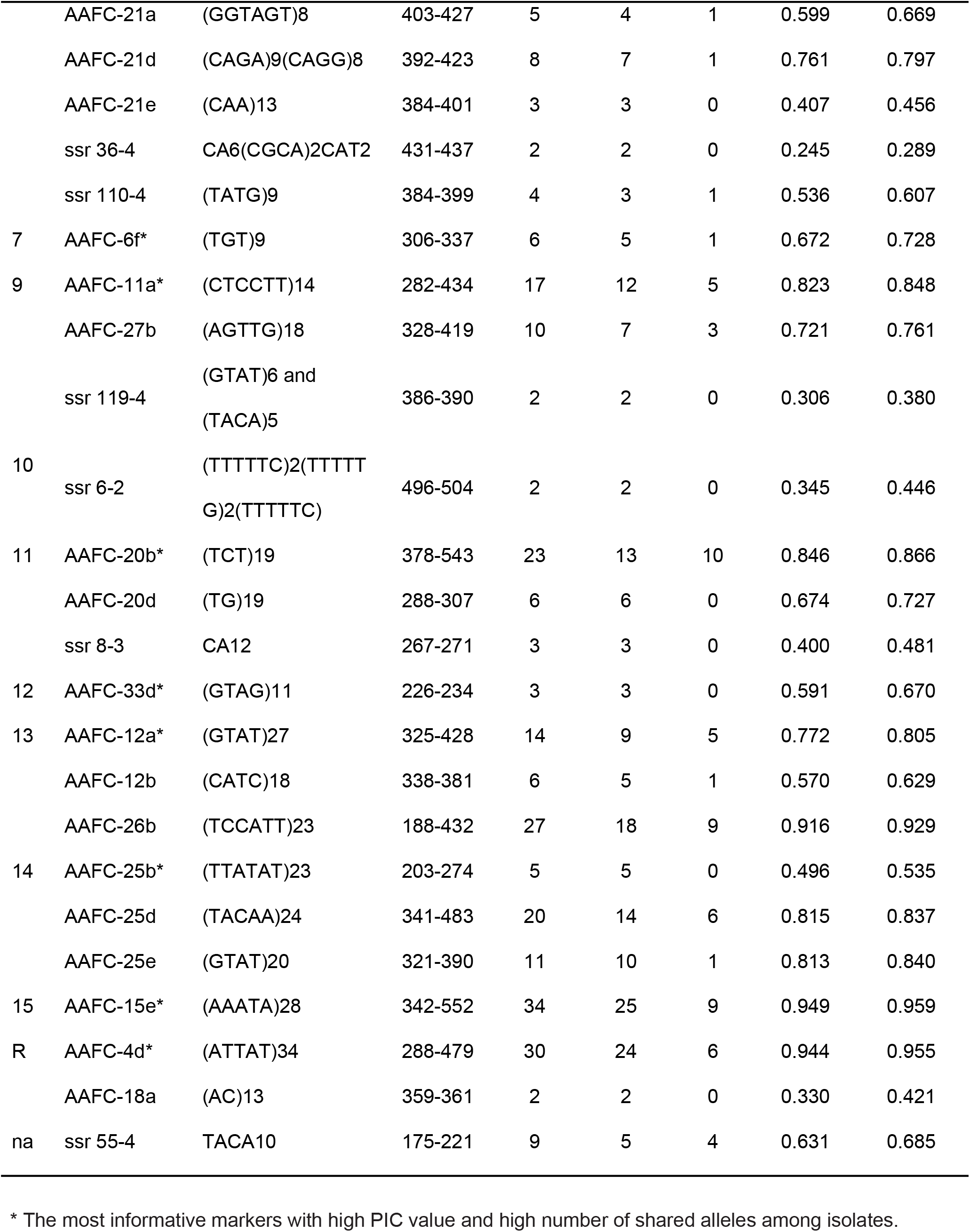
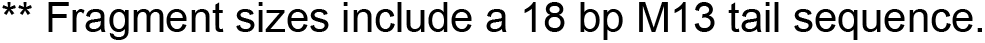
Information on simple sequence repeat (SSR) markers used for genotyping of *S. sclerotiorum* isolates organized by chromosome. Markers with AAFC-prefix were developed in this study, while markers with ssr-prefix were published by [4].

### Effect of geographic location

When SSR data from *S. sclerotiorum* isolates were grouped by province, analysis of molecular variance (AMOVA) showed 97% of the genomic variation was explained by differences among isolates, while 3% was due to differences among provinces (*P* = 0.001) (Table 3). Other analysis also showed the effect of geographical location. Isolates in Manitoba had the highest allelic richness (6.02) and the highest number of private alleles (1.25) compared to isolates in Saskatchewan and Alberta (Table 4). As expected, the genomic distance (*D*) between *S. sclerotiorum* populations was higher between the two distant provinces, Alberta and Manitoba (*D* = 0.098) (Fig 1), than between neighbouring provinces Saskatchewan and Manitoba (*D* = 0.017) and Saskatchewan and Alberta (*D* = 0.05) (Table 5). Congruently, gene flow (*Nm*) was highest between neighbouring provinces Manitoba and Saskatchewan (*Nm* = 53.97), followed by Saskatchewan and Alberta, and the lowest gene flow occurred between the two most distant provinces, Alberta and Manitoba. The differentiation index (*PhiPT*) was not significant between the neighbouring provinces Manitoba and Saskatchewan, but was significantly different between both Saskatchewan and Alberta, as well as between Manitoba and Alberta (Table 5).

**Table 3.** Analysis of molecular variance (AMOVA) based on simple sequence repeat markers in 127 *S. sclerotiorum* isolates from commercial canola fields in three Canadian provinces, Alberta, Saskatchewan and Manitoba (*P* = 0.001).

| Province | Allelic richness | Private allelic richness |
| --- | --- | --- |
| Manitoba | 6.02 ± 0.63 | 1.25 ± 0.22 |
| Saskatchewan | 5.66 ± 0.62 | 0.82 ± 0.17 |
| Alberta | 3.95 ± 0.32 | 0.85 ± 0.18 |

**Table 4.** Analysis of allelic richness among *S. sclerotiorum* isolates from three Canadian provinces.

| Source of variation | Df | Mean squares | Estimated variance | % of total variation |
| --- | --- | --- | --- | --- |
| Provinces | 2 | 35.3 | 0.5 | 3 |
| Isolates within Provinces | 126 | 15.3 | 15.3 | 97 |

**Table 5.**
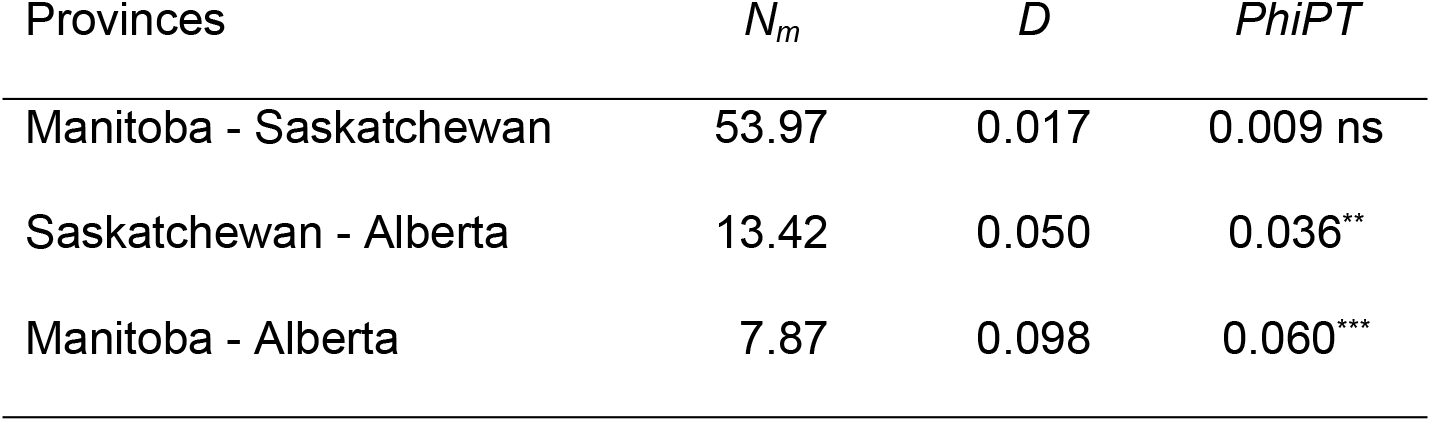
Analysis of gene flow (*Nm*), genetic distance (*D*) and population genetic differentiation (*PhiPT*) among *S. sclerotiorum* isolates collected in three Canadian provinces.

### Test of random recombination

Analysis of linkage disequilibrium in *S. sclerotiorum* assessed both by province and combined for the three provinces showed the Index of association (*I_A_*) was statistically significant in all cases, thereby rejecting the null hypothesis of random recombination. Also, all standard index of associations (*rBarD*) were much closer to 0 than to 1 specifying non-random association (Table 6). Thus, genomic variation based on SSR polymorphisms is less likely a result of random recombination in the ascospore stage, but rather through other mechanisms as discussed later.

**Table 6.** Analysis of linkage disequilibrium (*LD*) among *S. sclerotiorum* isolates from three Canadian provinces resulting in an index of association (*I*_*A*_) and a standardized index of association (*rBarD*).

| Province | $I_A$ | $rBarD$ |
| --- | --- | --- |
| Manitoba | 3.09** | 0.0685 |
| Saskatchewan | 4.36** | 0.0960 |
| Alberta | 2.78** | 0.0621 |
| Combined | 2.97** | 0.0654 |

### Population structure

Analysis of *S. sclerotiorum* population structure showed relatively high Delta K values supporting the existence of either 2, 12, 17 or 20 sub-populations (Fig 4). Existence of two sub-populations were highly significant with 63% of isolates in Q1 (25 isolates from AB, 28 from SK, and 28 from MB), 33% in Q2 (2 isolates from AB, 16 from SK, and 23 from MB), and 4% in an admix group (isolates MB18, MB27, MB24, MB35 and SK55) (S1 Table). Evidently, Saskatchewan and Manitoba isolates were almost equally represented in Q1 and Q2, but skewed towards Q1 in Alberta. Additional analysis of genomic distance among isolates visualized as a phylogenetic tree resulted in a multitude of possible sub-populations. Based on the results from these two types of analyses, it was decided 17 sub-populations best captured the genomic diversity, since Delta K was lower for 12 sub-populations, while 20 sub-populations did not add more clarity. The 17 sub-populations consisted of 1 to 22 isolates marked as alternate red and blue groups in Fig 5. One isolate from each sub-population was selected to represent the genomic diversity of the *S. sclerotiorum* population in western Canada, and they were subsequently evaluated for aggressiveness on *B. napus*.

**Fig 4.**
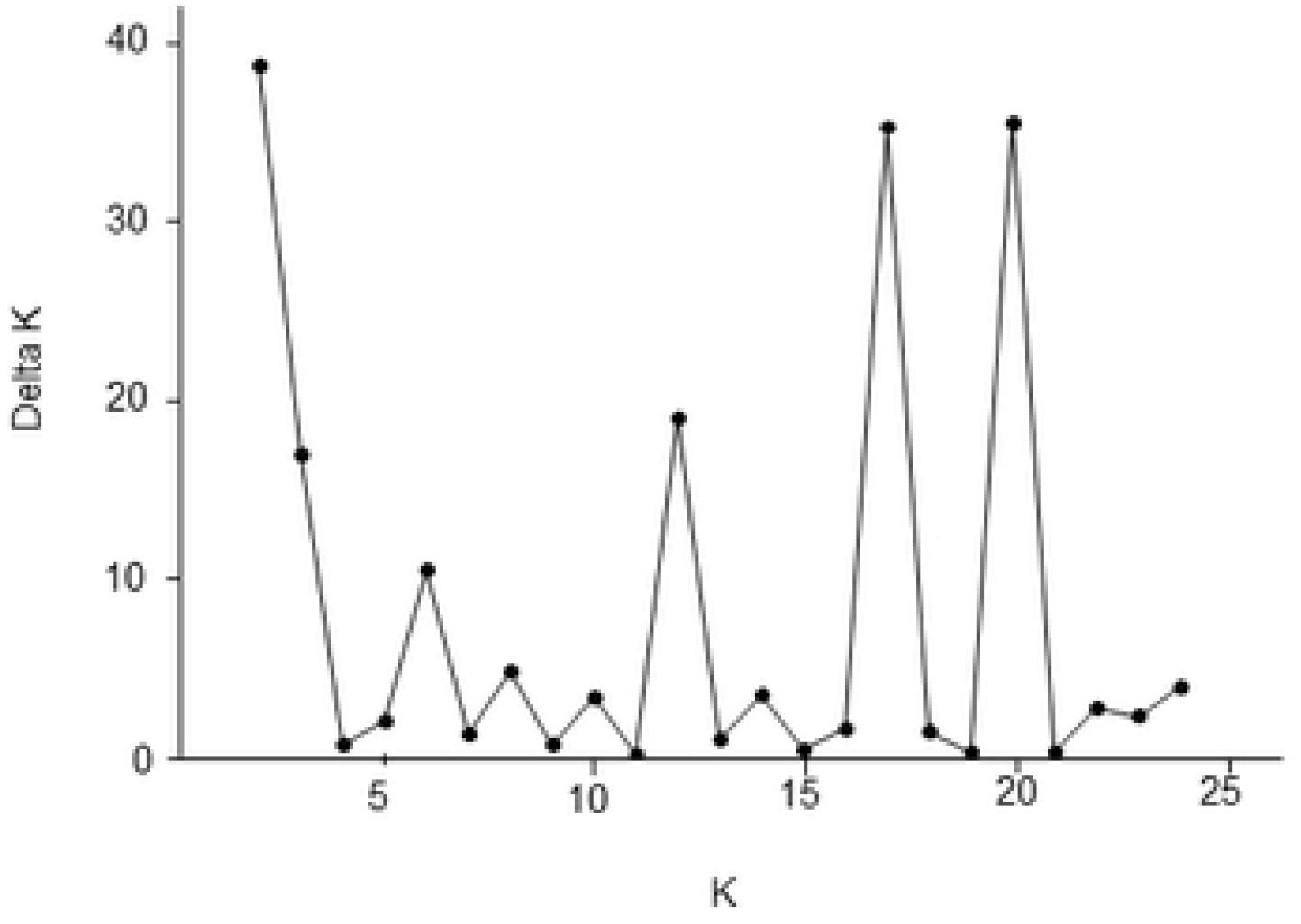
Likely number of sub-populations in *S. sclerotiorum* based on polymorphisms at 39 simple sequence repeat loci. The number of sub-populations among 127 isolates was determined using the Evanno method and graphed in Structure Harvester, a web-based program that visualizes output data from Structure (47).

**Fig 5.**
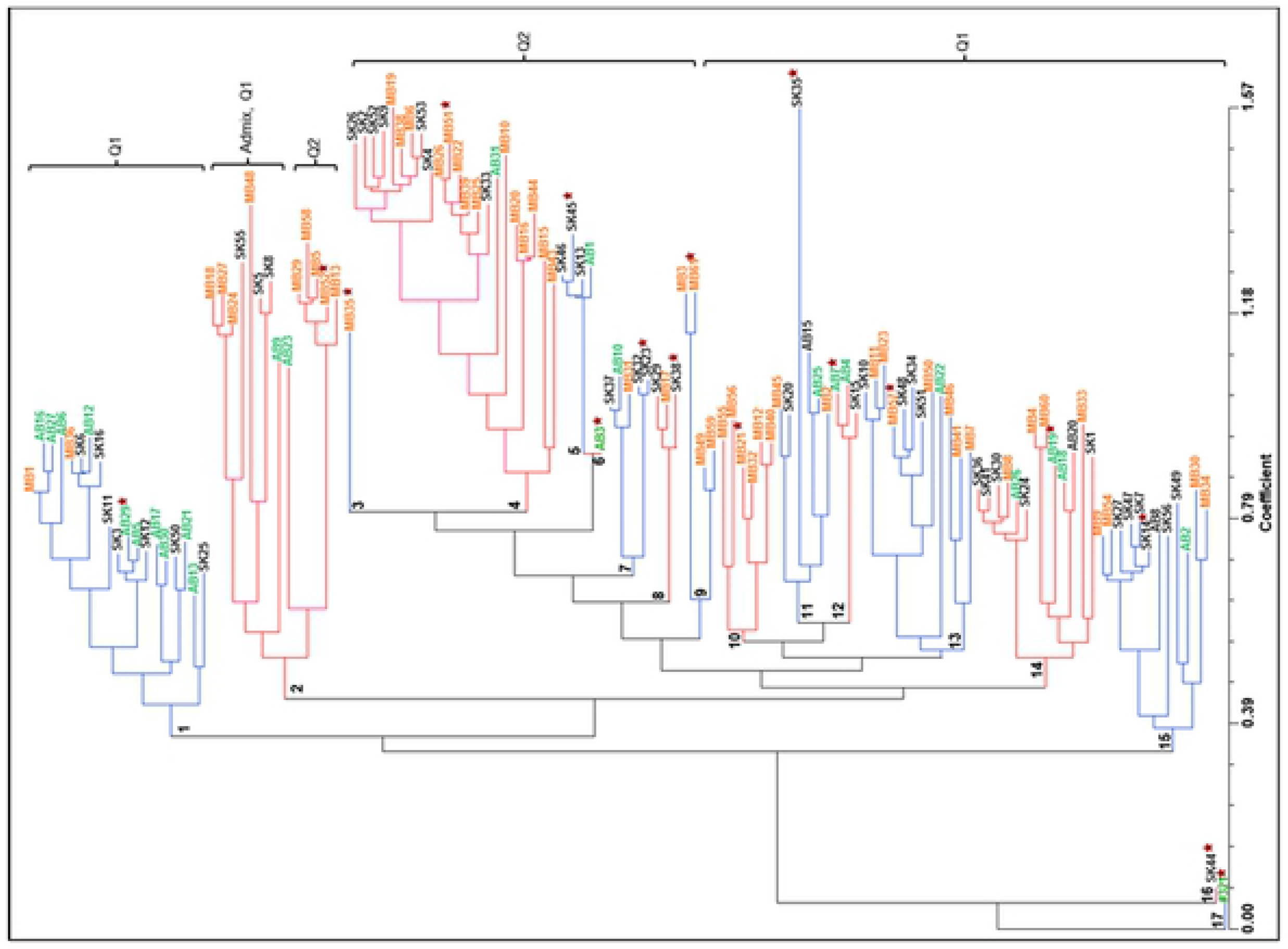
Phylogenetic tree of the relationship among 127 *S. sclerotiorum* isolates based on polymorphisms at 39 simple sequence repeat loci. Sub-populations are marked from 1 to 17 and coloured in alternate blue and red. The star shows one isolate from each sub-cluster selected for evaluation of aggressiveness.

### Isolate aggressiveness

The 17 *S. sclerotiorum* isolates were inoculated onto six *B. napus* lines separately. Disease progression was measured at weekly intervals as lengthwise colonisation of the stem and depth of penetration into the stem tissue measured as soft + collapsed lesions. Lesion length at each of three rating dates and the area under the disease progress curve (AUDPC) were highly correlated. Lengthwise lesion growth also was correlated with depth of penetration (S2 Table). Thus for simplicity, only the results from stem lesion length measured 21 days after inoculation are reported here. Analysis of variance (ANOVA) was significant for both *S. sclerotiorum* isolate and *B. napus* line. The lesion length for each isolate across six *B. napus* lines showed a continuum from the least aggressive isolate AB7 (17.4 + 3.2 mm) to the most aggressive isolate AB29 (151.3 + 13.8 mm) (S3 Table). Correspondingly, the lesion length on each *B. napus* line across 17 isolates ranged from the highest level of quantitative resistance in PAK54 (48.3 + 3.2 mm) to susceptibility in Topas (161.2 + 6.6 mm) (S4 Table). Lines could be divided into four groups based on LSD values with PAK54 most resistant followed by PAK93 and K22, then DC21 and Tanto. Interestingly, isolate by line interaction was significant (Table 7), which was particularly evident when stem lesion length for each *S. sclerotiorum* isolate was graphed for the six *B. napus* lines separately (Fig 6); this graph shows similar ranking of isolates from low to high aggressiveness on all *B. napus* lines, except isolate SK35, which was more aggressive on K22 and DC21 than on PAK54 and PAK93 (S5 Table).

**Table 7.** Analysis of variance of six *B. napus* lines, DC21, K22, PAK54, PAK93, Tanto and Topas, inoculation with 17 *S. sclerotiorum* isolates measured as stem lesion length 21 days after inoculation.

| Source of variation | Df | Mean squares | F-value | P-value |
| --- | --- | --- | --- | --- |
| *B. napus* lines | 5 | 107077.1 | 88.9 | 0.001 |
| *S. sclerotiorum* isolates | 16 | 25818.1 | 21.4 | 0.001 |
| Replications | 3 | 3664.8 | 3.04 | 0.029 |
| Lines x isolates | 80 | 2831.1 | 2.4 | 0.001 |

**Fig 6.**
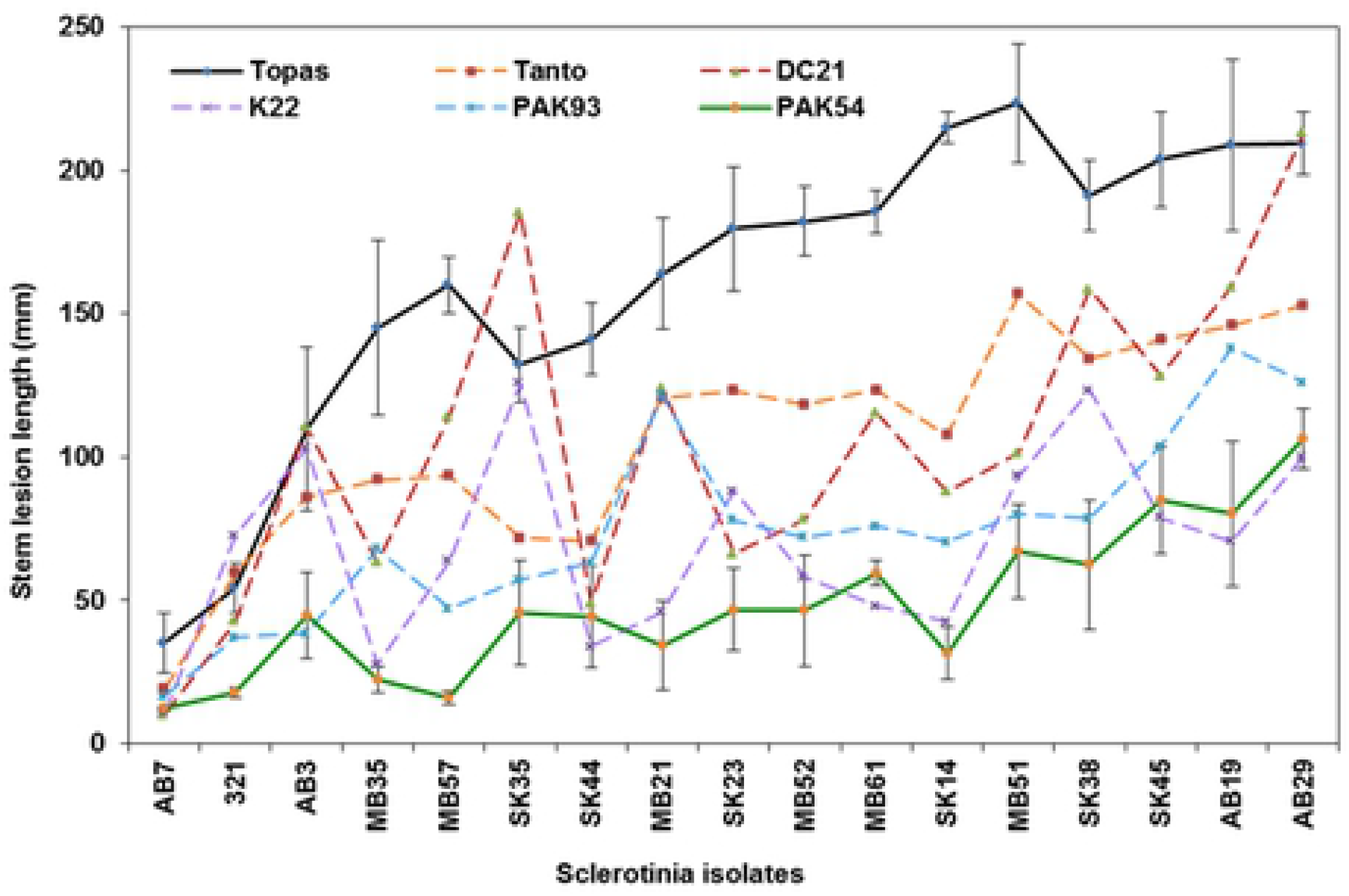
Results from evaluation of 17 *S. sclerotiorum* isolates for aggressiveness on six *B. napus* lines. Plants at full flower were inoculated by attaching a mycelium plug to the main stem with Parafilm. The average stem lesion length 21 days after inoculation is shown. For clarity, the standard error bar is only shown for susceptible Topas and resistant PAK54 (S5 has all standard errors).

## Discussion

Effective genotyping of *S. sclerotiorum* isolates from canola in a large geographical area combined with a new way of visualizing mycelium compatibility relationships gave us an informative ‘snap-shot’ of the pathogen population in western Canada. The sequenced *S. sclerotiorum* genome [13] was utilized to design SSR markers, and flourescent capillary electrophoresis allowed detection of single basepair size differences at 39 SSR loci distributed over the fungal genome. The resulting SSR polymorphisms were used to determine the relative contribution of isolates and geographical location to genomic diversity, linkage disequilibrium, population structure, and phylogenetic relationships among isolates.

*Sclerotinia sclerotiorum* has two nuclei in each ascospore and in cells of actively growing hyphal tips [14], while older and less organized mycelium contain myriads of nuclei. The two nuclei functions as a dikaryon for most of the pathogen’s life cycle, except for a brief phase during meiosis, when the 16 chromosomes condense into eight structures [15], providing an opportunity for genomic recombination. New allele combinations during meiosis are created by alignment of non-homologous chromosomes, followed by crossover events, whereby DNA strands break and re-join resulting in progenies with a genomic combination different from the parents. Since non-homologous recombination involve two genetically different nuclei, it occur at a low frequency in *S. sclerotiorum* for the following reasons; (1) meiosis takes place only once during the pathogen’s life cycle often corresponding to a single cropping season; (2) since the pathogen is homothallic, having both mating type genes at the same locus, it readily produce ascospores by self-fertilization, which out-competes other propagation scenarios; and (3) mycelium incompatibility prevent karyogamy between two different nuclei, while mycelium compatibility occur between genetically similar isolates.

Microconidia contain a single nucleus and a few organelles [16], and are found intermittently within mycelium and on the surface of sclerotia. Although the function of microconidia has not been determined, it is conceivably that these nuclei could transfer to hyphal tips during formation of asci, which takes place just below the melanised rind of the sclerotium [17]. In the event that nuclei in microconidia are genetically different from nuclei in the sclerotium, mixing of nuclei followed by non-homologous recombination is a slight possibility. Eskin [18] identified a morphological trait in a single *S. sclerotiorum* isolate having 5% of asci with four small and four larger ascospores. This size dimorphism might result from of mixing of different nuclei affecting some asci. However, in the present study, analysis of linkage disequilibrium rejected the hypothesis of random recombination in *S. sclerotiorum*, leading us to conclude non-homologous recombination is absent or extremely rare in the fungal population in western Canada.

Significant discoveries were made using diagrams visualizing the relationship among mycelium compatible isolates. Some *S. sclerotiorum* isolates were connected in strings where ‘X’ was compatible with ‘Y’, and ‘Y’ with ‘Z’, while ‘X’ was incompatible with ‘Z’, which resembled the ‘ring-species’ concept most often described for bird species (19). Most significantly, the five and six mutually compatible isolates in clone A and D (Figs 2 and 3) had almost identical SSR haplotype, and also belonged to a single sub-population 2 and 4, respectively (Fig 6) demonstrating they were closely related. Clones with fewer isolates, and isolates forming strings and pairs, belonged to several different sub-populations. In contrast, there were no similarity between SSR haplotype among the remaining incompatible isolates, which had numerous private alleles. The most likely sources of genomic diversity in *S. sclerotiorum* documented in this study consist of slippage during DNA replication and point mutation affecting individual nucleotides. Both mechanisms are particularly frequent in simple sequence repeats and accumulate with each mitotic cell division. The dataset seems to have captured isolates at various stages of divergence. Beginning with clonal isolates with similar SSR haplotypes, followed by stepwise divergence into compatible isolates forming strings, pairs of compatible isolates, and ending with incompatible isolates with unique SSR haplotypes. It is conceivable that genetic information passes from one mycelium compatible isolate to another by hyphal anastomosis, but over time, certain genetic factors prevent further compatibility, after which isolates become distinct haplotypes where polymorphisms continue to accumulate. In addition, it was clear that physical separation contributed to divergence, seen as low (11%) mycelium compatibility between isolates from different Provinces compared to 35-61% compatibility between isolates within Provinces.

Counts of shared and private alleles demonstrated each *S. sclerotiorum* isolates was a unique haplotype. This finding seems to contrast most previous publications, where haplotype frequencies were comparatively lower, but can be explained by the use of fewer SSR markers in those studies. Like us, these researchers used different sub-sets of markers published by Sirjusingh and Kohn in 2001 [4] to genotype *S. sclerotiorum* isolates from various plant species and geographic locations such as 6 SSR [20], 8 SSR [6, 21, 22, 23], 10 SSR [24], 11 SSR [25, 26], 12 SSR [27], and 13 SSR [28]. Understandably, using only 6 to 13 SSR markers limits the number of polymorphic alleles that can be detected, and consequently, pathogen populations are seen to comprise isolates with the same haplotype.

Although dividing isolates into Provinces seemed artificial at first, interpretation of mycelium compatibility and genomic diversity appeared to fit regional differences regarding crop rotation and weather conditions. Manitoba isolates had the highest allelic richness and private allelic richness, and also the highest proportion of mycelium compatible isolates (61%), all of which were related to one another in clones and strings (Fig 2). The fact that a high percentage of arable land in Manitoba is occupied by several susceptible crop species, canola, bean, soybean and sunflower (S6 Figure), compared to a lower percentage only occupied by canola in Alberta, together with higher frequency of wet weather conditions in Manitoba, has resulted in greater number of pathogen life cycles in both time and space, and therefore more opportunities for mycelium interaction among isolates in this province. Crop rotation and weather conditions in Saskatchewan fall between these two opposites. The results from the survey of *S. sclerotiorum* disease incidence in 2010 confirm these provincial differences (Table 1). The gene flow measured between provinces can be explained by planting of sclerotia-contaminated seed originating from other geographical areas, and infection from wind borne ascospores from distant fields both resulting in introduction of new haplotypes.

Analysis of population structure clearly divided the isolates into two sub-populations with 63% of isolates in Q1, 33% in Q2 and 4% in an admix group (S7 Table). Analysis of genetic distance visualized as a phylogenetic tree showed isolates could be further divided into 17 sub-populations. The two types of analysis were mostly in agreement, since the majority of isolates in each sub-population belonged to either Q1 or Q2 (Fig 5).

Genotyping of isolates collected from all canola producing areas of western Canada allowed selection of a practical number of genetically diverse isolates for subsequent evaluation of aggressiveness. Isolates ranked from low to high aggressiveness when inoculated onto a set of five *B. napus* lines with quantitative resistance, PAK54, PAK93, DC21, K22 and Tanto, based on stem symptoms which is the yield liming factor. Moreover, the isolate by line interaction was statistically significant, particularly evident for isolate SK35, which was more aggressive on DC21 or K22 than on PAK54 or PAK93 (Fig 6); interestingly, this isolate also had the most unique SSR haplotype (Fig 5). Similar specialization of *S. sclerotiorum* on host genotypes has been reported in several crop species including *B. napus* and *B. juncea* [5, 7], soybean [29], bean [30, 23], sunflower [31] and lentil [33]. Isolate MB51, collected in Lilyfield, Manitoba, and AB29, collected in Cayley, Alberta, were among the most aggressive isolates and also represented the two largest sub-populations of 22 and 19 isolates, respectively. These isolates would therefore be suitable for resistance screening during development of varieties destined for production in western Canada. The two *B. napus* lines, PAK54 and PAK93, which were partially resistant to a single isolate in a previous study [11], showed a high level of resistance against all isolates in the present study (Fig 6 and S5 Table). These lines showed quantitative resistance when evaluated against Australian isolates [33]. We are currently developing pre-breeding lines from PAK54 and PAK93 that combine sclerotinia resistance with good agronomic traits, and appropriate seed characteristics including high oil, low glucosinolate and erucic acid.

It is well known, *S. sclerotiorum* ascospores are unable to infect intact plant tissue, but first require uptake of nutrients from dead organic material in order to form infection cushions containing hundreds of hyphal tips, which in turn are capable of penetrating the plant’s epidermis [34]. Badet et al. [35] found *S. sclerotiorum* has undergone selective pressure toward optimization of a plethora of metabolites making it a generalist able to infect a wide range of dicot plant species, in contrast to specialized plant pathogens, such as *Zymoseptoria tritici* (causing septoria leaf blotch in wheat), which infects the host directly. Derbyshire et al. [36] concluded *S. sclerotiorum* has undergone a slow rate of evolution based on a low decay of linkage disequilibrium and a lack of selective sweeps in the genome, a process through which a new advantageous trait increases in a population that also can lead to reduced genetic variation in surrounding nucleotides. The latter investigation included five isolates, SK35, 321, MB52, MB21 and AB2, also part of the present study. Taken together, *S. sclerotiorum* is a relative weak and unspecialized pathogen that rely on secondary metabolites in the infection phase. It is therefore unlikely changes in aggressiveness in the pathogen population will overcome quantitative resistance in new varieties since the primary source of genomic variation is slippage during DNA replication and point mutation, while the probability of non-homologous recombination is low. Still, it is prudent to evaluate crop varieties against *S. sclerotiorum* isolates that are representative of the genomic and pathogenic diversity in the area where they will be planted.

## Materials and methods

### Collection of isolates and disease survey

In 2010, commercial canola fields were surveyed for the presence of *S. sclerotiorum* in all important canola producing areas of Alberta, Saskatchewan and Manitoba. A total of 168 fields were selected at random separated by at least 25 kilometers (Fig 1). The longitude and latitude of each location were recorded using a Global Positioning System (TomTom, Netherlands) (S7 Table). The incidence of *S. sclerotiorum* in each field was determined by counting the number of plants with typical stem rot lesions in a row of 10 plants at five sites (N = 50) separated by at least 10 meters (Table 1). Fields where only a few plants had lesions on leaves, side-branches or pods were rated as ‘trace’.

Sclerotinia-infected plants were present in 88% (149) of the fields. Isolates were made from 28 fields in Alberta, 53 in Saskatchewan and 55 in Manitoba for a total of 136 fields. In each field, infected stems were collected at ten sites separated by at least 10 meters for subsequent isolation of the pathogen. In addition, 200 infected stems were collected in a single field in Saskatchewan with 30% disease incidence. Four isolations were made from each plant, either from sclerotia in the stem pith or from infected stem tissue. The isolates were labelled with the acronym of the province (AB, SK or MB) a field number and the letters a, b, c or d. The sclerotia and stem pieces were surface-sterilized in 0.6% sodium hypochlorite for three minutes, rinsed in sterile water and plated on potato dextrose agar (PDA, Difco, Sigma-Aldrich, USA) in 9 cm Petri plates. Cultures were incubated in a cycle of 16 h day (22 +1°C) and 8 h night (18 +1°C), and after three to four days hyphal tips from the edge of a growing colony were transferred to a new PDA plate and incubated as before. Sclerotia that formed along the edge of the Petri plates were collected after four to six weeks and stored in paper envelopes under dark and dry conditions at 4°C with an identical set at −10°C. Examination of mycelium compatibility (MC), genotyping and testing for aggressiveness were conducted with isolates labeled ‘a’.

### Mycelium compatibility tests

Mycelium compatibility in *S. sclerotiorum* was examined using 133 isolates representing 28 fields in Alberta, 51 in Saskatchewan and 54 in Manitoba. Isolate 321 collected in 1992 from a canola field in Olds was part of the Alberta group [12]. In addition, 36 isolates were selected to represent a single, heavily infected field in Saskatchewan. Sclerotia of each isolate were surface sterilized, plated on PDA and incubated in a cycle of 16 h day (22+1°C) and 8 h night (18 +1°C). After 5-7 days, 4 mm plugs were cut from the growing margin of each culture. One mycelium plug of two different isolates were placed 3.5 cm apart in a 9 cm Petri plate on PDA supplemented with 75 μl/L McCormick’s red food coloring and incubated in the dark at 22+1°C as described by Schafer and Kohn [37]. Each isolate was paired with itself as a control of self-compatibility. In the first round, all isolates within each province, and those within the single, heavily infected field, were paired in all possible combinations for a total of n*(n-1)/2 pairings, where n is the number of isolates in each group. In the second round, mycelial compatibility was examined between provinces by paring 18 isolates against each other representing 3 fields in Alberta, 7 in Saskatchewan and 8 in Manitoba. All pairings in both the first and second rounds were carried out twice. A compatible interaction showed continuous mycelium growth over the entire Petri plate. In contrast, a incompatible interaction showed a barrage zone of sparse mycelium between the two isolates often with a red line in the media, that was particularly evident on the reverse side of the Petri plate. Plates were examined visually after 7 and 14 days and scored as + or - on the day the interaction type was most evident. Initially, the data were entered in a traditional isolate by isolate scoring matrix (S2 Table). Subsequently, diagrams were created to visualize all compatible interactions by province, inter-provinces and in a single field as shown in Fig 2 and 3.

### Genotyping

Simple sequence repeats were identified in the sequenced *S. sclerotiorum* genome available on the Broad Institute’s web site [13]. A total of 32 SSRs were selected to represent 15 chromosomes and one contig predicted in this assembly and given the prefix AAFC (Table S8). Primer pairs for PCR amplification of these SSRs were designed using WebSat software [38]. In addition, 15 primer pairs for amplification of other *S. sclerotiorum* SSRs were obtained from Sirjusingh and Kohn [4] and given the prefix ssr (Table S8). Genotyping was carried out with *S. sclerotiorum* isolates from group ‘a’ described above. In preparation for extraction of genomic DNA, sclerotia of each isolate was surface sterilized, cut in half and placed on PDA in a 9 cm Petri plate. After 5-7 days incubation at 16 h light (22+1°C) and 8 h dark (18 +1°C) two 4 mm plugs were cut from the growing margin and transferred to potato dextrose broth in a 9 cm Petri plate and incubated as before. When mycelium covered 80% of the liquid surface it was harvested, washed twice with sterilized, distilled water and lyophilized.

Total genomic DNA was extracted from 30 mg ground mycelium using a DNA isolation kit according to the manufacturer’s protocol (Norgen Biotek Corp, ON, Canada). DNA was quantified using a Quant-it PicoGreen Assay (Invitrogen, USA) on an Appliskan microplate reader (Thermo Fisher Scientific, USA) and diluted to 10 ng DNA μl^−1^. Each PCR reaction was performed in a 8.2 μl total volume containing 2.6 μl @ 10 ng μl^−1^ fungal DNA template, 0.0625 μM M13 primer fluorescently labeled with one of FAM, VIC, NED, PET or LIZ (Applied Bio-Systems), 0.67 μM of each of forward (with M13 tail) and reverse primer, and 4.1 μl of FideliTaq PCR master mix (FideliTaq 2x, 25 mM MgCl_2_, 20 mM dNTPs). PCR run conditions were optimized for BioRad DNA Engine Dyad (BIO-RAD, USA) resulting in the following amplification conditions: 94°C for 3 min, then 22 cycles at 94°C for 30 s, 56°C for 30 s, 72°C for 45 s, followed by additional 22 cycles of at 94°C for 10 s, 56°C for 20 s, 72°C for 1 min; with a final extension of 10 min at 72°C. PCR reactions for each dye were pooled, and 3 μl of each pooled aliquot were mixed with 7 μl formamide plus size standards, denatured at 95°C for 5 min, and ice chilled. Fluorescent capillary electrophoresis was run on an ABI 3730xl DNA Analyzer (Applied Biosystems, USA).

Initially, the quality of each primer pair was examined by gel electrophoresis of PCR amplification products from a sub-set of eight *S. sclerotiorum* isolates (data not shown). Eight SSRs were eliminated due to poor performance: ssr 7-2, ssr 7-3, ssr 17-3, ssr 20-3, ssr 110-4, ssr 114-4, AAFC-2b and AAFC-2d. Subsequently, 30 AAFC and 9 ssr primers were used to genotype 127 *S. sclerotiorum* isolates. Fragment size analyses were performed using Genographer 2.1.4 [39]. Size differences of one or more base pairs were considered separate SSR alleles. The size estimation at hyper variable loci with more than 20 different alleles were carried out manually. All possible alleles were determined for each SSR locus and a matrix was generated indicating the presence (1), absence (0) or no amplification (null) of each allele in all 127 isolates. Only unambiguous data with >80% amplification success rate were included in the final data set.

### Analysis of genomic diversity

The Genographer dataset was curated into Excel (Microsoft, USA). Allele counts, including total number of polymorphic alleles, shared and private alleles, were conducted in the Microsatellite feature of the Toolkit program [40]. The polymorphic information content (PIC) value for each primer pair across isolates was calculated using the GenAIEEx 6.5 feature in Excel [41]. Genomic diversity (*H*) was calculated in POPGENE 1.32 [42]. The haplotype was determined for individual isolates combining alleles across all loci using FaBox 1.41 [43] (data not shown).

The SSR data were grouped into the three provinces, Alberta, Saskatchewan and Manitoba, where the isolates were collected and used for several types of analyses. Contribution of isolate and geographical location to genomic variation was determined by analysis of molecular variance (AMOVA) using GenAlex v 6.5 with 999 permutations, and the results presented in Table 3. Analysis of allelic richness and private allele richness among isolates within each province was carried out using the software ADZE 1.0 with the rarefraction approach where sample size was equal to the smallest sample size [44] (Table 4). Analyses of gene flow (*Nm*), Nei’s unbiased genetic distance (*D*), population differentiation (*PhiPT*) for pairwise comparison of provinces were calculated using GenAlex 6.5 in Excel with 999 permutations [41] (Table 5). Two analyses of linkage disequilibrium (*LD*) were used to test the null hypothesis that random recombination exists. For this, the index of association (*I*_*A*_) and the standardized index of association (*rBarD*) were calculated using the software Multilocus v.1.31 [45] for each province and the three provinces combined (Table 6). The null hypothesis was tested by comparing expected and observed values with 1000 permutations; so that if *rBarD* equal 0 there was a non-random association of alleles, whereas *rBarD* equal 1 specify random association of alleles.

### Examination of population structure

A Bayesian cluster analysis was used to infer genomic ancestry among the 127 *S. sclerotiorum* isolates using Structure [46]. Analyses with the number of sub-populations ranging from K = 2 to K = 24 were performed using Markov Chain Monte Carlo replication method. The admixture ancestry and independent allele frequency options were used throughout. The number of sub-populations among isolates was determined using the Evanno method and *ln(P)* was graphed in Structure Harvester, a web-based program that visualizes output data from Structure [47]. The resulting Delta K values are graphed in Fig 4.

The relative genomic distance among *S. sclerotiorum* isolates based on SSR polymorphisms was analyzed using NTSYSpc 2.2 with matrices of genetic distance coefficient function ‘SIMGEND’ sub-program with Nei72 [48]. Subsequently, neighbour-joining analysis (Njoin) was performed with default parameters. To visualize the results, the ‘TREE’ function was used to generate a phylogenetic tree, where the oldest isolate, 321 collected in 1992, was selected as the root and branches consisted of all other isolates collected in 2010 (Fig 5).

### Evaluation of aggressiveness

Based on analyses of population structure (Delta K) and the phylogenetic tree, 17 *S. sclerotiorum* sub-populations were selected to represent the genomic diversity in western Canada. One isolate from each sub-population was selected for evaluation of aggressiveness so that each province was represented by five to six isolates as follows: Alberta isolates AB3, AB7, AB19, AB29 and 321; Saskatchewan isolates SK14, SK23, SK35, SK38, SK44 and SK45; and Manitoba isolates MB21, MB35, MB51, MB52, MB57 and MB61. The isolates were inoculated onto six *B. napus* lines that were selected based on their phenotypic reaction to a single *S. sclerotiorum* isolate, 321 [11]. These lines were PAK54 and PAK93 (Pakistan), DC21 (South Korea), K22 (Japan), with high level of quantitative resistance, variety Tanto (France) with intermediate resistance and variety Topas (Sweden) as a susceptible control.

Seeds of each line were sown into water-soaked peat pellets (Jiffy-7, McKenzie, Brandon, MB, Canada) and placed in a greenhouse. Plants at the 3-4 leaf stage were transplanted into natural soil in a phenotyping facility under semi-field conditions. The facility consisted of a 20 m x 40 m greenhouse structure with retractable roof and side walls made from reinforced polyethylene supported by permanent gable ends of corrugated polyvinyl (Cravo, Brantford, ON, Canada). Plant growth relied on ambient sun light and temperature with rainfall supplement with overhead irrigation when needed. A weather sensor and computer program operated motors that closed the roof and side walls to protect the plants at temperatures below 10°C and wind speeds above 30 km per hour. At temperatures above 25°C the plants were cooled by activation of an overhead misting system and shaded by closing the roof 50%.

Stocks of the 17 isolates were produced by inoculating the susceptible canola variety Topas with each isolate and re-isolating the pathogen onto nutrient rich V8 juice media (200 ml V8 juice, 0.75 g CaCO3, 800 ml water, 15.0 g agar). After 4-6 days, 7 mm plugs of mycelium were transferred to cryo-freezer solution (10% skim milk, 40% glycerol) and stored at −80°C until needed for inoculation. Inoculum of individual isolates was prepared by transferring mycelium plugs from the cryo-freezer onto PDA plates. The plates were incubated in a cycle of 16 h day (22 +1°C) and 8 h night (18 +1°C), and were ready for inoculation after 4-5 days at which point the culture was still actively growing, but had not reached the edge of the Petri plate. Mycelium plugs were cut with a 7 mm cork borer from the margin of actively growing cultures and placed on 3 × 7 cm pieces of stretched Parafilm (Bemis Company Inc, Oshkosh, WI, USA) with the mycelium facing up. When each plant was at full flower, two internodes of the main stem were inoculated by attaching a mycelium plug with Parafilm as described by Gyawali et al. (11). The length of the developing lesions was measured at 7, 14 and 21 days after inoculation (dai), and subsequently used to calculate the area under the disease progress curve (AUDPC). In addition, depth of penetration into the stem tissue was assessed as either firm, soft or collapsed and used to calculate percent soft + collapsed lesions for each isolate and line. The experiment consisted of six *B. napus* lines in four replications planted as the main plot, with the 17 *S. sclerotiorum* isolates inoculated onto plants as sub-plots organized in a randomized complete block design (RCBD). There were seven plants in each sub-plot for a total of 2,856 plants. The mean values for each combination of *S. sclerotiorum* isolate and *B. napus* line were calculated for the five disease traits (7 dai, 14 dai, 21 dai, AUDPC and % soft + collapsed lesions).

Correlations between these traits were analysed in a pairwise manner using the Pearson correlation coefficient (PROC CORR) in SAS Enterprise 5.1. The result showed all traits were highly correlated (S2 Table). For simplicity, only stem lesion length measured at 21 dai are presented here. Analysis of variance (ANOVA) was conducted using the PROC GLM model in SAS Enterprise 5.1 to determine variation explained by isolate, *B. napus* line, and the interaction between isolate and *B. napus* line (Table 7), followed by calculation of the least significance difference (LSD) between entries (S3 and S4 Tables).

## Acknowledgement

Creation of the map of *S. sclerotiorum* sample locations by Cam G Kenny is greatly appreciated.

## Supporting Information

**S1 Table.** Mycelial compatibility scoring matrix by Province, inter-Province and single field.

**S2 to S5 Tables.** Correlation among disease traits in tests of aggressiveness. Average aggressiveness by isolate. Average aggressiveness by line. Standard Error for all comparisons.

**S6 Fig.** Map of canola production areas in western Canada.

**S7 Table.** *Sclerotinia sclerotiorum* sample locations, isolate names, Q1 and Q2 values from population structure analysis.

## Author Contributions

LB conceived and lead the research and wrote the manuscript. LB and DDH secured funding. AD collected isolates. HG designed primers and amplified SSRs. AS and KKG sequenced SSR products. KDP and JD performed statistical analysis. JA, MH, DL, AD and HG performed tests of mycelium compatibility and aggressiveness. All authors reviewed the manuscript.

